# Tracking of quiescence in *Leishmania* by quantifying the expression of GFP in the ribosomal DNA locus

**DOI:** 10.1101/641290

**Authors:** Marlene Jara, Ilse Maes, Hideo Imamura, Malgorzata A. Domagalska, Jean Claude Dujardin, Jorge Arevalo

## Abstract

Under stressful conditions some microorganisms adopt a reversible non or slow proliferative quiescent stage that allows their survival. Although quiescence has been described broadly in bacteria, this phenotype has been only recently discovered in *Leishmania*. In the present work we developed a biosensor of quiescence that allows to monitor the physiological stage of the parasite at population and single cell levels. We inserted a GFP gene into the ribosomal DNA locus and followed the expression of this reporter gene, driven by the ribosomal promotor (rGFP expression). We showed that rGFP expression decreased significantly and rapidly during the *in vitro* transition from extracellular promastigotes to intracellular amastigotes of *L. mexicana* an *L. braziliensis* and that the decrease in rGFP expression was coupled *in vitro* with a decrease in replication as measured by BrdU incorporation. Quiescence was not only observed in reference laboratory strains, but also among clinical isolates. We found that quiescence was reversible as the parasites could rapidly resume their metabolically active and proliferative stage when they were put back in an optimal environment for growth. We demonstrated for the first time in live cells that amastigotes are a heterogeneous population in which shallow and deep quiescent stages may coexist. Finally, we showed that rGFP expression could be monitored *in vivo* and that quiescent amastigotes could reside in tissues of animals with latent infections of *L. braziliensis* or *L. mexicana*. We propose rGFP expression as a simple parameter to define quiescent cells and further characterize them.

**IMPORTANCE:** Quiescence is a physiological diversification that allows pathogens to overcome chemotherapy without the development of drug resistance and to be invisible to the immune system of their host. Quiescent pathogens can cause latent infections and (re-) emerge in an unpredictable time during the lifetime of the individual. The phenomenon was recently described in *Leishmania* in which it could explain several clinical and sub-clinical features, like therapeutic failure, reactivation of the disease and asymptomatic infections. However, a simple biosensor of quiescence for *Leishmania* is not yet available. We show for the first time that the integration of GFP within the rDNA locus and the subsequent quantification of its expression can be used as a biosensor to distinguish quiescent subpopulations among live amastigotes. Moreover, we show quiescence is quickly reversible both *in vivo* and *in vitro*. We offer a tool that will allow the further molecular characterization of quiescent parasites.

## INTRODUCTION

*Leishmania* is a digenetic parasite involving two life stages. The motile promastigotes reside in the invertebrate sand fly and the non-motile amastigotes reside within the parasitophorous vacuole of the macrophage and other phagocytic cells (1–3). In the mammalian host the parasite can produce a broad spectrum of clinical manifestations ranging from local skin lesions known as cutaneous leishmaniasis, to metastasis and severe mutilations in oropharyngeal tract known as mucosal leishmaniasis and to a systemic form, known as visceral leishmaniasis, which is fatal in the absence of treatment. One common feature of all infections is the sub clinical persistence of a low number of parasites after therapy or even decades after clinical cure (4–7). These kinds of infections resemble the well known and characterized latent infections of *Mycobacterium tuberculosis*, where the pathogen remains within its host in a physiologically quiescent stage for long periods, if not the lifetime of the individual. Quiescence is a metabolically downregulated stage with no or slow proliferation and it is considered to be the metabolic status of pathogens in latent infections (8, 9). Under optimal conditions *in vitro M. tuberculosis* has a generation time of 18-24 hrs, while the time of generation in latent infections *in vivo* rises to ~100 hrs (10, 11). Although pathogens can persist indefinitely in a quiescent stage, these latent infections represent a threat for their host, as pathogens can also resume their fast growth and produce the recrudescence of the disease in an unpredictable time during the lifetime of the individual.

Although there is clinical evidence of the persistent condition of *Leishmania* infections, there are scarce studies about quiescence in *Leishmania*. In 2015, Kloehn *et al.* used for the first time the term quiescent to characterize the metabolic stage of amastigotes in *L. mexicana in vivo*. Analyzing the whole population of cells, they found several quiescence traits among amastigotes: a substantially increased generation time in comparison to promastigotes (12 days vs 9 hours, respectively), plus lower synthesis of RNA, proteins, lipids (12). Our group achieved the same conclusion for another species, *L. braziliensis* and we described a drop in the levels of macromolecules involved in the ribosome biogenesis (ribosomal RNA and proteins (13), a signature of quiescence encountered from prokaryotes as *Mycobacterium* to eukaryotes as *Sacharomyces* (14–16). Quiescence complexity was further addressed by a third study that explored the homogeneity of quiescence among *L. major* amastigotes *in vivo (17)*. The authors found that in persistent infections of the self-healing C57BL/6J mice model, there are two populations among amastigotes: one in a slow proliferative stage (generation time of 60 hours, semi quiescent) and another with no evidence of proliferation (quiescent). These results suggested that amastigotes can thus adopt different physiological stages during the course of the infection.

Molecular mechanism triggering quiescence in *Leishmania* are still unknown and specific molecular markers are not yet available. As the amastigotes may present two physiologically different stages, one quiescent but another proliferative, there is a need for a biosensor that would allow to distinguish both sub-populations for further molecular characterization. Considering that transcription of rDNA is (i) regulated by a well-defined promotor in *Leishmania* -in contrast to other genes- and (ii) down-regulated in quiescent stages, we hypothesized that the expression of a reporter gene integrated in the rDNA locus could serve as a biosensor of quiescence. We used here GFP as a reporter gene and demonstrated in *L. braziliensis* and *L. mexicana* reference strains that it can be used as a biosensor of the proliferative stage of parasites and as a tool to distinguish quiescent stages among amastigotes at a single cell level. Using this biosensor, we have characterized *in vitro* the heterogeneity of physiologically downregulated stages among *Leishmania* amastigotes and the kinetic of the entrance and exit of these stages. The results of experimental work *in vivo* and the evaluation of quiescence in a panel of clinical isolates are also shown.

## RESULTS

### rGFP expression in *Leishmania* promastigotes and amastigotes

The decreased transcription of the rDNA locus is a trait of quiescence in several microorganisms (15, 20, 21). Therefore, we postulated that if the GFP gene was integrated within the rDNA locus, the subsequent measurement of its expression (rGFP expression) would serve as a quiescence biosensor and would allow to discriminate quiescent populations among alive parasites. In *L. braziliensis*, the analysis at whole population level showed that rGFP expression in alive cells (as indicated by their integrity of membrane with NucRed Dead staining) was the highest in the highly proliferative Pro^Log^, with an average value of 1995.0 ± 5.7 (mean RFU ± SD). The rGFP expression decreased progressively and significantly to 1361.0 ± 6.2, 217.3 ± 46.6 and 23.37 ± 2.6 in Pro^Sta^, Amas^Axe^ and Amas^Int^ respectively (One way ANOVA, *P*= 0.0002) (Fig 1 A-C). In *L. mexicana* the same sequential and significant decrease in rGFP expression was observed between the same life stages. (Fig S1).

**Fig 1.**
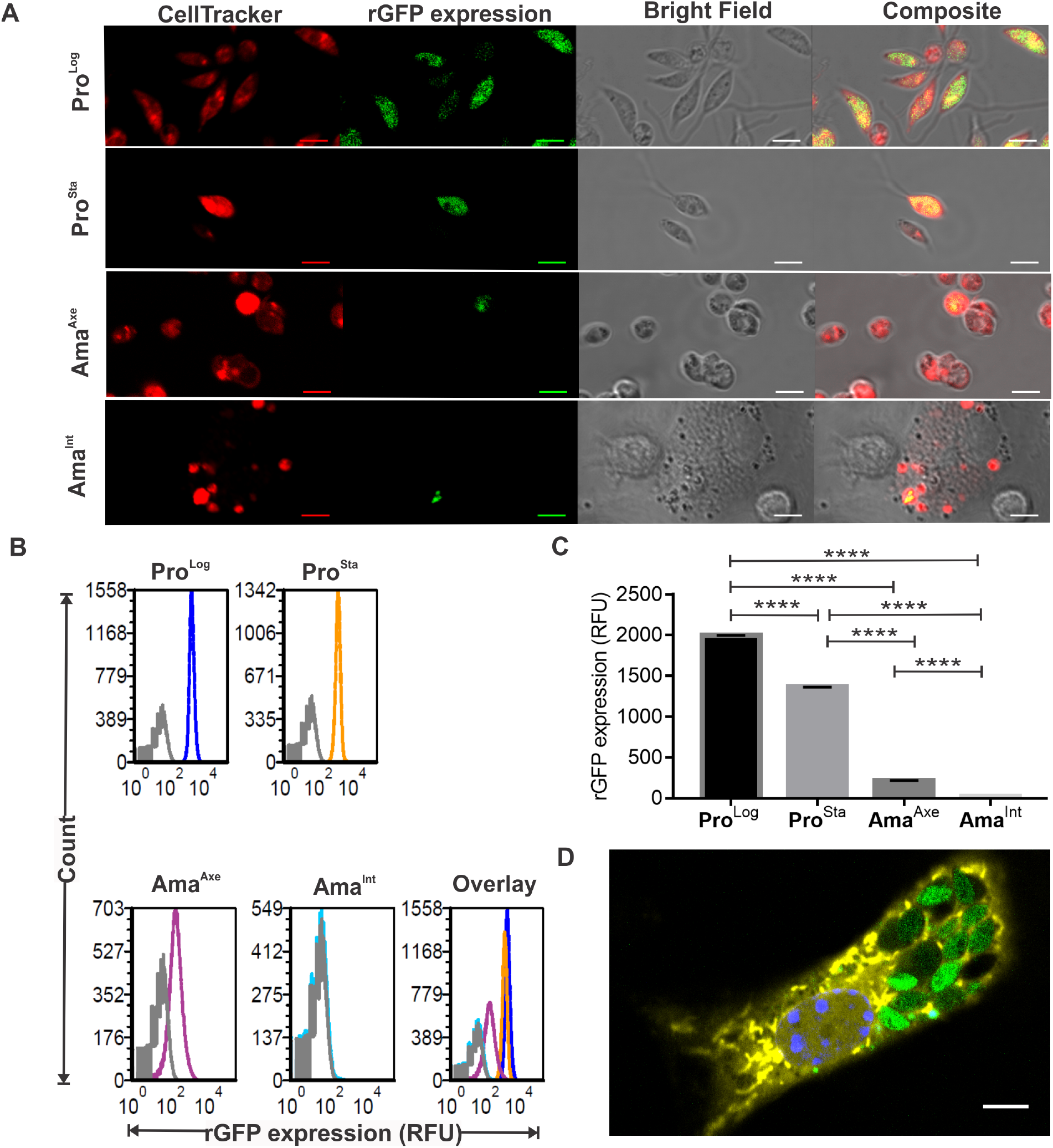
*Leishmania* amastigotes have a downregulated rGFP expression and constitute a more heterogeneous population in comparison to promastigotes. **A**, Live cell imaging of *L. braziliensis* M2904 rGFP Cl2. The expression of GFP integrated into the ribosomal locus (rGFP expression) was monitored by confocal microscopy in Pro^Log^, Pro^Sta^, Ama^Axe^ and Ama^Int^ (respectively logarithmic and stationary promastigotes and axenic and intracellular amastigotes). The Cell tracker Deep Red was used to detect each parasite cell. **B** and **C**, evaluation of rGFP expression over the cell cycle of *L. braziliensis* in alive cells (negative for NucRed Dead fluorescence) by flow cytometry. **B**, Analysis at single cell level of the heterogeneity of rGFP expression among promastigotes and amastigotes: the broader the peaks, the more heterogeneous the cell population. The gray peak in each panel is an overlay of the same life stage in the wild type strain (Non GFP negative control). Each peak represents 10^5^ live cells. **C**, Median of rGFP expression in each life stage (measure is made at population level). The mean of three biological replicates is shown. **D**, Macrophage infected with different physiologically downregulated stages of *L. mexicana* amastigotes. The scale bar in each figure represents 5 μM.

We further analyzed the flow cytometry results at single cell level. We observed that for both species, Ama^Axe^ constitute a more heterogeneous population than promastigotes, as evidenced by the thickness of the rGFP expression histogram (Fig 1 B). The analysis of the variances of each population in *L. braziliensis* showed that Ama^Axe^ have statistically significant higher variability (coefficient of variation (CV)= 90,2) in comparison to Pro^Log^ (CV=36,0) or Pro^Sta^ (CV=36,9) (F test < 0.0001). The acquisition of new images with settings to optimize the detection of GFP confirmed the heterogeneity of downregulated stages in Ama^Int^ even among the ones residing in the same macrophages (Fig 1D). We then calculated in each life stage the proportion of cells that have quantifiable levels of rGFP expression. Using a very stringent gate which allows only 0.04% of false positive in the negative control (non rGFP *Leishmania* strain), we found that in *L. braziliensis* the percentage of cells with quantifiable rGFP expression was 99.6 ± 0.1, 99.4 ± 0.1, 52.3 ± 12.8 and 0.56 ± 0.2 (mean ± SD) in Pro^Log^, Pro^Sta^, Ama^Axe^ and Ama^Int^ respectively (Fig 2A). Very similar results were shown in *L. mexicana*: in Ama^Int^ of that species, 4.0 ± 0.7 % (mean ± SD) reached quantifiable but low rGFP expression (Fig 2B). This population can be observed in the Fig S1B as a small peak at the right side of the corresponding graph (arrow). Overall, the results show that the downregulation of rGFP expression is an inherent trait of amastigotes and that it develops rapidly as it is already seen in Ama^Axe^.

**Fig 2.**
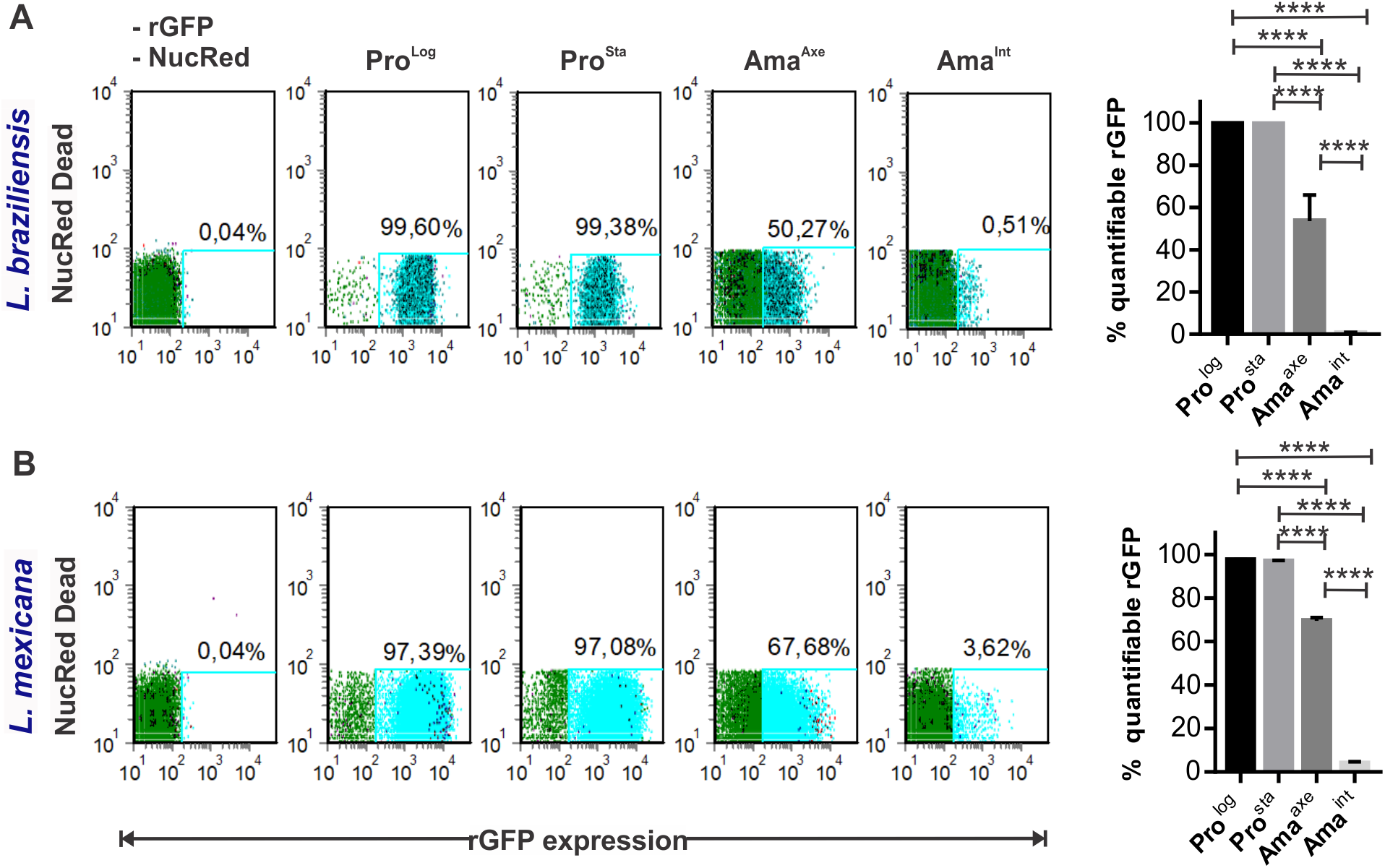
Evaluation of the proportion of viable cells with quantifiable expression of rGFP in promastigotes and amastigotes of *L. braziliensis* (A) and *L. mexicana* (B), by flow cytometry. Each scatter plot of one biological replicate represents the rGFP expression (X-axis) vs NucRed Dead fluorescence (Y-axis), negative in live cells. The light blue gate represents the population with cell viability and quantifiable expression of GFP. The histograms on the right represent the % of the population with quantifiable rGFP expression (Mean of 3 biological replicates +/− SD). The asterisks represent samples with statistically significant difference (One way Anova *P*< 0.0001).

### Evaluation of rGFP expression and its relationship with cell proliferation

The deeply downregulated levels of rGFP expression in amastigotes indicated that amastigotes are in a quiescent stage which is characterized by a low or non-proliferative condition. In order to establish the relationship between rGFP expression and the proliferative condition of parasites we evaluated simultaneously both parameter by immunofluorescence. We quantified (i) the proportion of proliferative cells with the use of the BrdU assay which is the standard molecular method to measure DNA replication, together with (ii) the immunofluorescence of rGFP. Images were acquired with a confocal microscope and the quantification of both parameters was performed by flow cytometry (Fig 3). Fast incorporation of BrdU and high rGFP expression (~ 80000 RFU) was observed in Pro^Log^ under conditions where they were replicating with a doubling time of ~ 8 hrs. After 4, 8 and 24 hrs of incubation with BrdU, there was 47, 80 and 90 % of BrdU^+^ Pro^Log^ cells respectively. As expected, with the decreased rGFP in Pro^Sta^ (~ 30000 RFU) most of the cells were BrdU ^−^ with only a small proportion of the population (~ 10%.) being BrdU^+^ for all the times of exposition. In both promastigote stages of *L. braziliensis* the mean of rGFP expression on the population BrdU^+^ was significantly higher in comparison to the one of BrdU^−^ (Fig. 3D).

**Fig 3.**
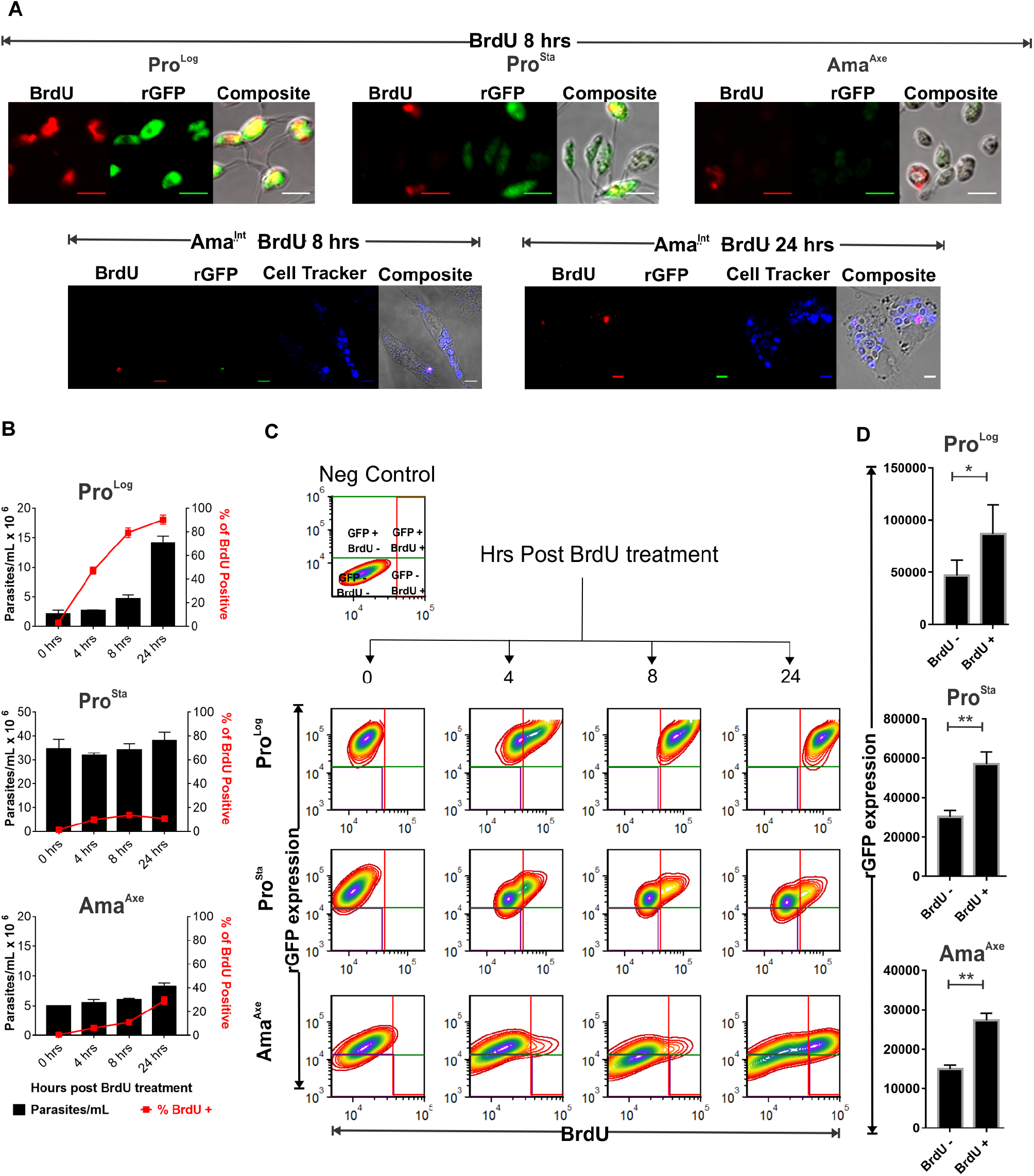
Relationship between BrdU incorporation (as a measure of replication) and rGFP expression in promastigotes and amastigotes of *L. braziliensis*. **A**, Confocal microscopy pictures of the immuno-detection of GFP and BrdU, 8 hrs after exposure of the cells to BrdU. The images for Pro^Log^, Pro^Sta^ and Ama^Axe^ were taken with the same microscopical settings to allow a comparison between the samples. The images for Ama^Int^ were taken with increased settings of the Gain Master to have a higher sensitivity for the detection of GFP. The new settings became the rGFP expression detectable in a small proportion of them. The scale bar in each picture represents 5 μM. **B**, Quantification by flow cytometry of the percentage of cells in a proliferative stage as evidenced by the incorporation of BrdU. The bars and the dots represent the mean ± SEM of three biological replicates. **C**, Density plot of rGFP expression (Y-axis) and level of BrdU incorporation, after flow cytometry. **D**, Quantification of rGFP expression in proliferative (BrdU^+^) and non-proliferative (BrdU^−^) populations of each stage, after 8 hrs of exposure to BrdU. The bars represent the mean ± SEM of three biological replicates.

The population of Ama^Axe^ did not show cellular duplication over 24 hrs, which is in agreement with their even more decreased levels of rGFP expression (~ 16000 RFU) and slow although progressive increase in their BrdU^+^ population, evincing their diminished rate of proliferation. After 4, 8 and 24 hrs of incubation with BrdU, there was 6, 11 and 29 % of BrdU^+^ cells respectively. Taking into consideration the incorporation of BrdU after 4 hrs of incubation, we calculated that the replication time in *L. braziliensis* Pro^Log^ is ~ 7.6 hrs while it would be ~ 62.4 hrs in Ama^Axe^ if we consider them as a homogeneous population. However, considering the results above, the occurrence of two physiologically different sub populations in Ama^Axe^ is likely: one with delayed incorporation of BrdU and low but positive rGFP expression and a second one with negligible incorporation of BrdU and not quantifiable rGFP expression (Fig 3C). In this case the adjustment of the generation time for the rGFP^+^ population gives a replication time of 48.8 hrs. In the rGFP^−^ cells, generation time (if there is any replication) cannot be estimated.

### Evaluation of the reversibility of the quiescent stage

We evaluated the kinetics and reversibility of quiescence by the quantification of rGFP expression. The arrested or slowly growing population should be able to revert to its faster rate of growth once parasites are in an environment without nutritional limitations or stressful conditions. With this purpose we analyzed the rGFP expression in *L. braziliensis* and *L. mexicana* daily during the process of differentiation from promastigotes to Ama^Axe^ (in response to the acidic pH and high temperatures). In *L. braziliensis* the levels of rGFP expression during the process of amastigogenesis in comparison to Pro^Log^ decreased significantly to 64 ± 0.2 % (mean ± SD), 43.1 ± 0.8 %, 28.2 ± 1.4 % and 19.8 ± 0.1 % on day1, day 2, day 3 and day 4 respectively (One way ANOVA, P<0.0001) (Fig 4A). Similar results were found during the amastigogenesis of *L. mexicana*: rGFP expression decreased to 88.5 ± 2.2 (mean ± SD), 38.9 ± 1.0, 18.7 ± 0.6 and 13.5 ± 0.9 % on day 1, day 2, day 3 and day 4 respectively (One way ANOVA P=0.003) (Fig 4A). We also evaluated the reversibility of rGFP expression when the parasites differentiate back from Ama^Axe^ to promastigote. The rGFP expression on the whole population of *L. braziliensis* increased progressively from day 1 (32.1 %) to day 3 (86.3 %) and growth was resumed after day 1. More heterogeneity was observed in *L. mexicana*: on day 1 we saw that although most of the population initiated the increase of their levels of rGFP expression, a fraction of them maintained a low rGFP expression as evidenced by the tail in the left of the histogram (Fig 4B).

**Fig 4.**
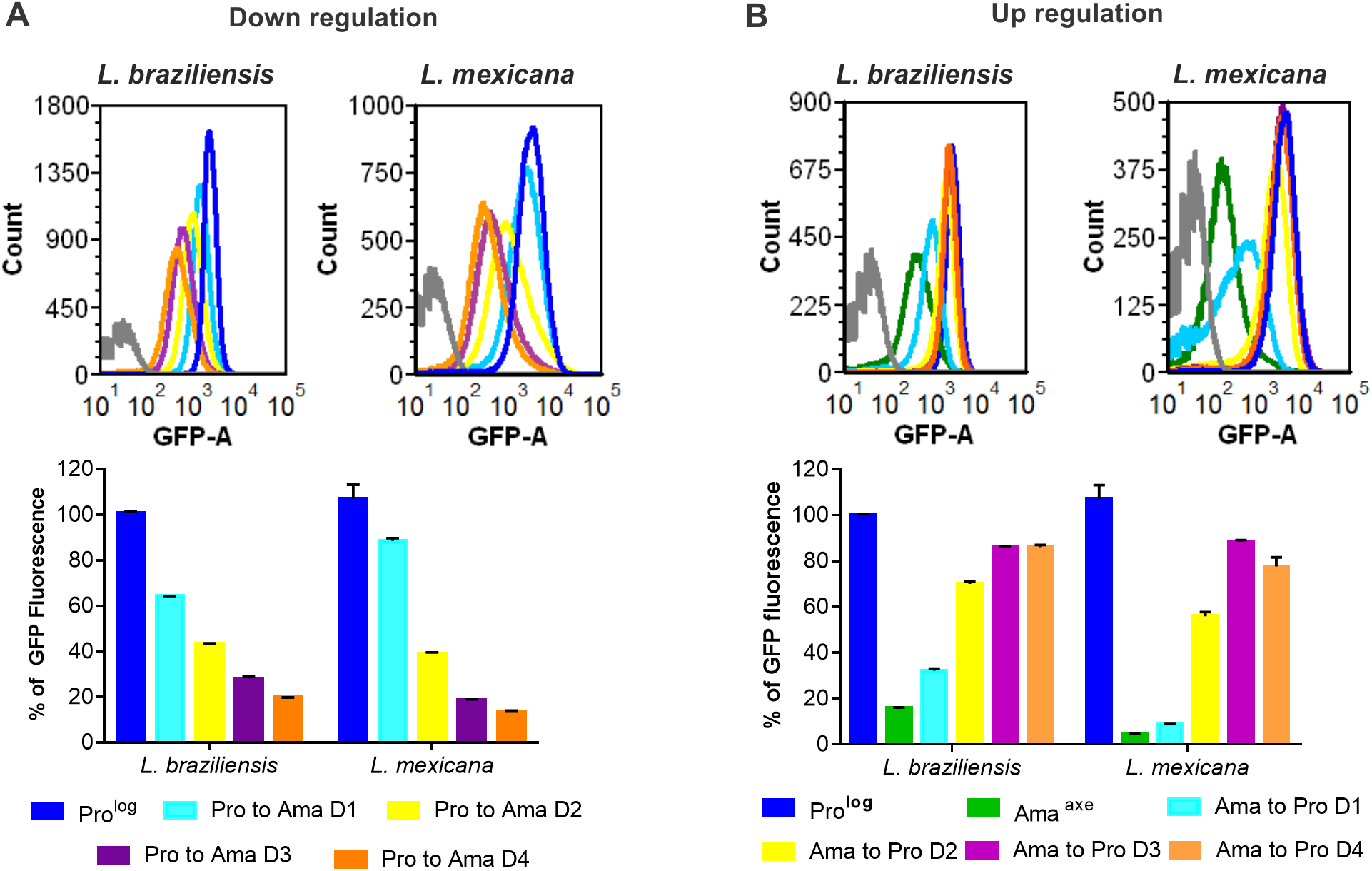
Kinetics of rGFP expression during amastigogenesis of *L. braziliensis and L. mexicana* (**A**) and during back-transformation from amastigotes to promastigotes (**B**). The expression of rGFP was quantified by flow cytometry. The grey peak in each panel represents the auto fluorescence of the non rGFP wild type line. The bars represent the mean ± SEM of three biological replicates.

### Evaluation of the quiescent stage of *L. braziliensis* and *L. mexicana* amastigotes in animal models

We further wanted to validate the use of rGFP expression as a marker of quiescence by monitoring amastigotes obtained from tissues with latent infections *in vivo*. If so, we were expecting that amastigotes *in vivo* would also have deeply downregulated levels of rGFP expression. In order to be able to track the quiescent amastigotes *in vivo* we performed the infection of animals with CellTracker Deep Red pre-stained promastigotes of rGFP lines. This staining should be useful to detect the amastigotes in case the rGFP was downregulated below the detection level. In addition, it could be used to confirm the quiescent condition of amastigotes. Indeed, proliferative cells are expected to dilute progressively the CellTracker Deep Red until it becomes non-detectable: this was verified in proliferative promastigotes, in which the dye becomes not detectable on day 6 (Fig 5 A-B). In contrast, in both hamsters infected with *L. braziliensis* and mice infected with *L. mexicana*, intracellular amastigotes positives for CellTracker Deep Red were visualized in all points of time (up to 120 days post infection). This suggests a very slow rate of proliferation. Also, the retention of CellTracker Deep Red correlated with the non-detectable (most of the events) or negligible rGFP expression in each visualized amastigote and amastigotes with high rGFP expression and no signal for the CellTracker Deep Red were never detected (Fig 5C). In order to confirm that the amastigotes with low rGFP expression do represent viable cells, an aliquot of the harvested cells was placed in Tobie’s medium to allow the differentiation of amastigotes back to promastigotes and their further propagation. For both species, promastigotes were recovered in 100% of the animals 60 days post infection. At 120 days post infection, promastigotes were recovered in 75% and 100 % of the animals in *L. braziliensis* and *L. mexicana* respectively (Fig 5 D). In order to confirm that the negligible expression of rGFP of *in vivo* amastigotes is due to their quiescent stage and not due to the loss of GFP within the rDNA locus, we measured the rGFP expression in the isolates recovered 60 days and 120 days post infection. All promastigotes differentiated from the mice-derived amastigotes recovered a pattern of rGFP expression comparable to the original parasites before the infection. For instance, the parasites isolated from 4 mice, 60 days post infection, generated populations with 99.2 ± 0.4 % (mean ± SD) of GFP positive cells and with 2077 ± 341 RFU (Fig 5E). Similar results were obtained with *L. mexicana*.

**Fig 5.**
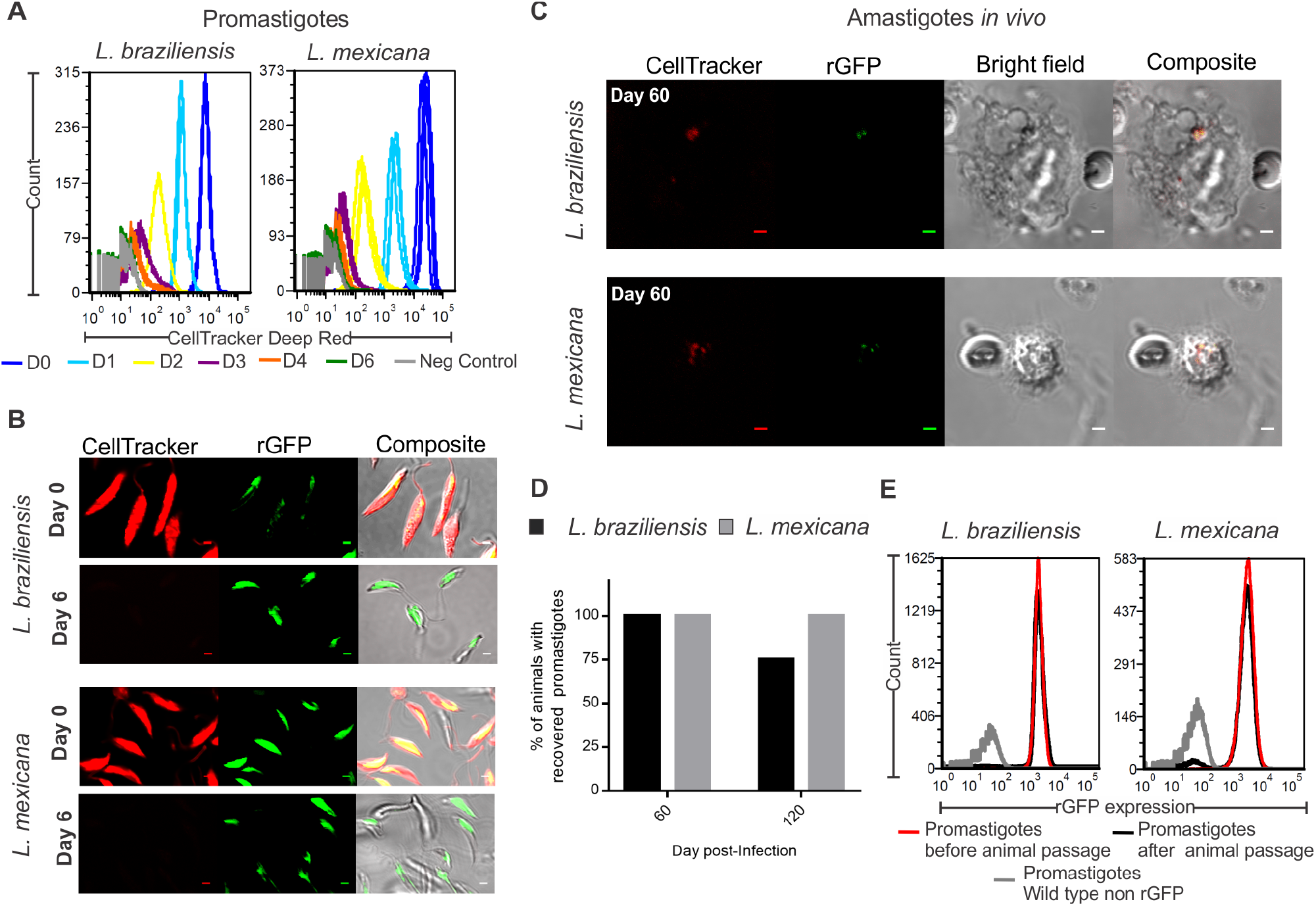
Flow cytometry (**A**) and confocal microscopy (**B**) of promastigotes stained with Celltracker Deep Red. Logarithmic promastigotes have a high expression of rGFP and in each cycle of replication the CellTracker is diluted within the cell. The CellTracker becomes non-detectable 6 days after staining. **C-E**, rGFP expression, proliferation, cell viability and reversibility of quiescence parameters in Ama^Int^ derived from hamsters (*L. brazilensis*) and mice (*L. mexicana*), 60 days post infection. The images for Ama^Int^ were taken with an increased setting of the Gain Master to have a higher sensitivity for the detection of GFP amastigotes. The scale bar in each picture represents 2 μM.

### Quiescence of *L. braziliensis* amastigotes in clinical isolates

In order to extend the relevance of our study to (recent) clinical isolates, we derived rGFP cloned lines from each of them and evaluated their capability to adopt a quiescent stage by measuring the rGFP expression (Fig. 6). We found that as observed in *L. braziliensis* M2904 rGFP Cl2, Ama^Axe^ of the 8 lines showed downregulation of rGFP expression in Ama^Axe^, with levels ranging from ~26 % to only ~6% of the levels of Pro^Log^ (Fig 6B). These results show all the clinical isolates can adopt a quiescent stage, and that there is some variability in the extend of downregulation among these.

**Fig 6.**
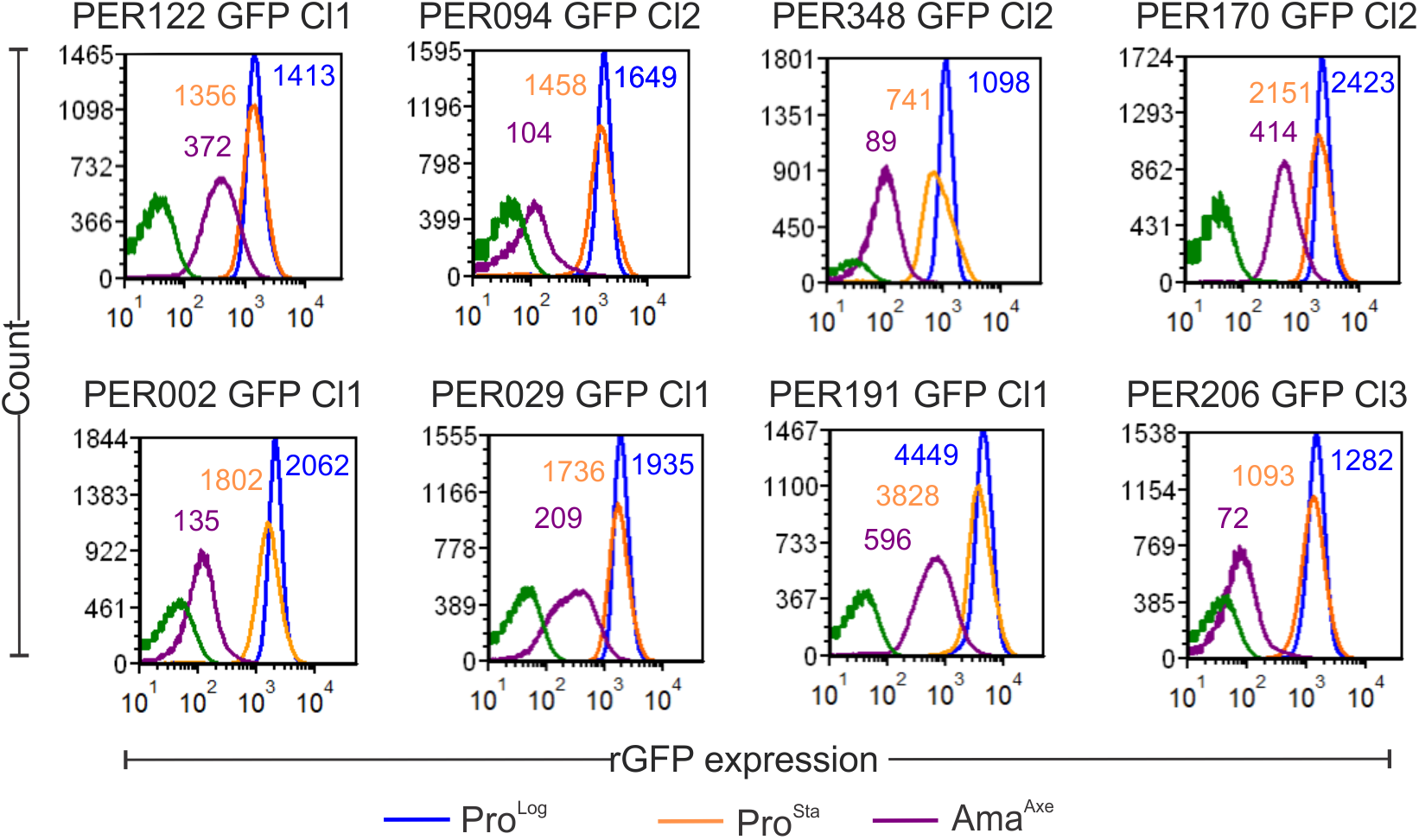
Quiescence in amastigotes is a common phenotype that can be found in clinical isolates of *L. braziliensis*. Overlays of the quantification of rGFP expression by Flow cytometry. The numbers in each peak indicates the mean of rGFP expression in three biological replicates.

## DISCUSSION

Many pathogens can remain in their host and cause a subclinical infection after chemotherapy or self-healing. These latent infections can extend during the lifespan of the host and constitute a reservoir for reactivation of the disease in an unpredictable time. This is the case in leishmaniasis, where relapse is common in the absence of drug resistance (22, 23), and complications like muco-cutaneous leishmaniasis or post kala-azar dermal leishmaniasis occur months to years after healing of primary cutaneous or visceral lesions, respectively (24, 25). In other pathogens, *in vivo* models of subclinical infections have shown evidence that hosts harbor the pathogen in a quiescent stage with no or slow replication (8, 9, 11, 26). However, the physiology of these stages is barely characterized *in vivo* because of their scarce number and the difficulty to detect them. We aimed here to develop and validate a simple bio-marker of quiescence and an *in vitro* model for further studies of this phenomenon in *Leishmania*. In this study, we inserted the GFP gene into the rDNA locus to evaluate if the expression of this reporter gene driven by the ribosomal promoter may serve as a biosensor of the metabolic activity of the parasite. We showed that (i) rGFP expression decreased significantly and rapidly during the *in vitro* transition from extracellular promastigotes to intracellular amastigotes, (ii) the decrease in rGFP expression was coupled *in vitro* with a decrease in replication as measured by BrdU incorporation, (iii) modulation of rGFP expression both *in vitro* and *in vivo* was reversible, the signal of the reporter gene increasing again during the transition from amastigote to promastigote and (iv) the use of rGFP expression allowed to identify heterogeneity among amastigotes and (v) our rGFP bio-sensor was applicable *in vivo*,.

A common feature of quiescent populations in a broad range of microbes is the reduction in the abundance of ribosomal RNAs and corresponding proteins (15, 27, 28). Down-regulation of biosynthetic machinery, including ribosomes, fits with cells that do not need to duplicate their DNA and synthetize all the proteins and other macromolecules to generate daughter cells. This was clearly demonstrated in *Mycobacterium bovis* where the fast proliferative stages had ~3880 ribosomes per cell while the slow proliferative stages had 687 ribosomes (29). In *L. infantum* Ama^Axe^ showed a decreased rate of synthesis of proteins and amount of polyribosomes (30). Similarly, our previous study in *L. braziliensis* showed that Ama^Axe^ downregulate their levels of rRNA to 15 % of the levels found in Pro^Log^ (13). In present study, we showed that Ama^Axe^ of both *L. mexicana* and *L. braziliensis* had an overall downregulated rGFP expression equivalent to 10% of the levels found in Pro^Log^. This reduction becomes even more stringent in Ama^Int^ as the overall expression of the rGFP dropped to less than 1% in *L. mexicana* and 3% in *L. braziliensis* in comparison to the levels found in Pro^Log^. This higher drop can be an adaptative mechanism that allows the amastigotes to survive in the phagolysosome where they will face the limitation of micronutrients and the macrophage microbicidal processes (31–33). We further extended the documentation of the quiescence phenomenon to clinical isolates. Considerable variability in the downregulation of rGFP expression suggests that clinical isolates could have different propensity to adopt the quiescent stage.

The inverse relationship between proliferation and biogenesis of ribosomes offers a set of potential molecular markers for identification of quiescent cells (34). In the present study, the results of the BrdU assay for proliferation confirm that rGFP expression can be used as a marker for the identification of quiescent populations of amastigotes as in Ama^Axe^ only the population with measurable but low rGFP expression showed some incorporation of BrdU while the population with no rGFP expression did not show any detectable incorporation of BrdU. The deep down regulation of biosynthesis (here measured by rGFP expression) and the negligible incorporation of BrdU in Ama^Int^ are in agreement with the constant infection index in peritoneal macrophages infected with *L. braziliensis* and *L. mexicana* over 72 hrs post infection: this was an early indicator of the lower rate of proliferation of Ama^Int^ in comparison to the ~ 7 hrs required by promastigotes (35–37). The use of rGFP as a marker of quiescence will facilitate further molecular characterization of quiescent subpopulations.

In pathogens reversibility of the quiescent stage to a proliferative condition is imperative to enable transmission and survival of the population (38). This was demonstrated here, as both axenic and rodent *ex vivo* amastigotes were also capable to rapidly resume their active biosynthetic level and proliferative condition when they were placed in conditions optimal to differentiate back to the proliferative promastigote stage.

*Leishmania* amastigotes may show a range of physiologic heterogeneity. It has been shown that 4 months after the infection with *L. major*, the healed foot pad of C57BL/6J mice harbors two populations of amastigotes: one that replicates with a time of generation of 60 hrs and a second one that do not show evidence of replication (17). Monitoring rGFP expression in *L. mexicana and L. braziliensis* amastigotes supported this concept of heterogeneity among amastigotes through the co-existence of amastigotes with varying downregulated physiological stages. Similarly, in *L. amazonensis* it has been shown that clones derived of parasites freshly taken from lesion have three phenotypes: fast, slow and intermediate progression to pathology in BALB/c mice (39). These phenotypes were not strictly genetically based as a fast clone generated sub clones with the three phenotypes (39). Our results in *Leishmania* cloned populations support the growing evidence that quiescence in pathogens is a dynamic phenomenon where subpopulations of shallow to deep quiescent stages may form the whole population (9, 40, 41). It is postulated that this heterogeneity allows the survival of the population rather than individual cells in different environmental conditions (38, 39).

*In vivo Leishmania* infections have different outcome depending on the strain, model and the number of parasites inoculated (38). In the present study we used *Leishmania* strains which upon inoculation generated asymptomatic infection during the 4 months of the experimental work. Although no lesions were developed, we found amastigotes in the tissues. The quiescent condition of parasites in these latent infections was confirmed by two indicators: i) the negligible expression of rGFP expression and ii) the retention of the CellTracker Deep Red up to 120 days post infection. Our results show that the inoculation of a moderate dose of stationary parasites in hamsters and mice for *L. braziliensis* and *L. mexicana* respectively (and the strains here evaluated) represents a good model for the further molecular characterization of quiescent population in latent infections. Although, the integration of fluorescent or luminescent reporters within the rDNA locus is useful to monitor quiescence, its application to monitor infections *in vivo* could be limited as the downregulation in the transcription of the rDNA locus may bring the expression of the reporter under the limit of detection of the *in vivo* imaging systems. Therefore, the lack of signal of the quiescence reporter gene does not necessarily indicate the absence of infection. A second reporter gene introduced in another locus, which transcription is constitutive or amastigote-specific should be developed as positive control.

We showed that *L. braziliensis* and *L. mexicana* strains produce a heterogeneous population of quiescent amastigotes that are characterized by a decreased rate of proliferation and a lower capacity of biosynthesis, both reversible. The use of transgenic rGFP parasites and single cell approaches will offer valuable tools for the further molecular *in vitro* studies of quiescent subpopulation among amastigotes. The relevance of quiescence in subclinical infections and its potential role in non-heritable drug tolerance in *Leishmania* urge additional exploration as it has been shown that it plays an important role in other pathogens (42–44).

## MATERIALS AND METHODS

### Ethics statement

Mice and hamsters were used for infections *in vitro* and *in vivo* by methods approved by the ethical committee from the Institute of Tropical Medicine (Antwerp). They certified that Protocols N° MPU2018-1 and MPU2018-2 followed the guidelines of the Federation of European Laboratory Animal Science

### Parasites strains and culture

Clones of *L. braziliensis* MHOM/BR/75/M2904 and *L. mexicana* MHOM/BZ/82/BEL21 were used in most of the experiments. If not otherwise indicated, the results showed in this manuscript correspond to these strains. We extended our findings to clones derived from 8 clinical isolates of *L. braziliensis* that were used for the characterization of rGFP expression over the cell cycle: MHOM/PE/03/PER191, MHOM/PE/03/PER206, MHOM/PE/02/PER029, MHOM/PE/01/PER002, MHOM/PE/02/PER122, MHOM/PE/04/PER348, MHOM/PE/04/PER170 and MHOM/PE/02/PER094 (provided by the Institute of Tropical Medicine Alexander von Humboldt). Promastigotes were cultivated in complete M199 (M199 medium with Glucose 5g/L to a final pH of 7.2 and supplemented with 20 % Fetal Bovine Serum (FBS), hemin 5 mg/L, 100 units/mL of penicillin and 100 μg/mL of streptomycin). In the next paragraphs the abbreviation Pro^Log^ and Pro^Sta^ will be used for logarithmic and stationary promastigotes, respectively.

In order to obtain axenic amastigotes, promastigotes (1.0 ×10^6^ parasites/mL) were maintained in complete M199. On the fourth day, 1 mL of promastigotes was centrifuged (1500 g) and the pellet was re-suspended in 5 mL of complete M199 at pH 5.5 and incubated at 34 °C. After four days, the resulting axenic amastigotes (Ama^Axe^) were harvested or used to obtain *in vitro* intracellular amastigotes (Ama^Int^) by the infections of BALB/c peritoneal macrophages at a ratio of 4 Ama^axe^ to 1 macrophage as previously described elsewhere (18). Three biological replicates per each stage and condition were evaluated in all the tests *in vitro*.

### Development of GFP *Leishmania* lines

GFP was integrated within the 18S ribosomal DNA locus with the use of the pLEXSY-neo2 system (Jena Bioscience) as previously reported elsewhere (19). After the generation of the transgenic rGFP parasites, they were cloned by the ‘micro-drop’ method. Briefly, an appropriate dilution of a 2 days old promastigote culture was prepared from which micro-drops of 1 μl were placed in the middle of the well (96 wells plate). In each well the presence of a drop with a single promastigote was verified microscopically and upon confirmation 150 μl of complete M199 medium was added. After incubation at 26°C for one week, growth was microscopically examined and established clones were transferred to a biphasic Tobie’s medium for further routine culture. The number of integrations of GFP within the rDNA loci was confirmed by whole genome sequencing of a PCR free library with a HiSeq 4000 platform (BGI). The sequences were assembled having as reference the genome of the strain M2904 as previously described elsewhere (18). The sequence analysis showed the presence of 1 and 2 copies of GFP per genome in the lines of *L. braziliensis* and *L. mexicana* respectively.

### Monitoring rGFP expression in living parasites by flow cytometry

Promastigotes and amastigotes of rGFP lines were stained for the evaluation of their cell viability by using the NucRed dead 647 (Thermo fisher scientific): labeling marks dead cells. The samples were analyzed with a calibrated flow cytometer BD FACS Verse™ in the medium flow rate mode. A wild type (non rGFP) line, not stained with NucRed was included in each experiment as negative control. In order to compare the fluorescence among samples the relative fluorescence units (RFU) were acquired with the same settings during all the experiments. The FCS files were analyzed with the FCS 5 express Plus research edition. Briefly, single cells were selected by pulse geometry gate; a first gate was selected with SSC-W and SSC-H, a second gate was created with FCS-H and FCS-A. Among single cells, a third gate was created by plotting the rGFP expression and the NucRed staining; the levels of rGFP expression was evaluated in cells negatives for the NucRed dead staining.

### Double immunodetection of BrdU and rGFP for confocal microscopy and flow cytometry

To test BrdU labeling in promastigotes and amastigotes the cells were cultured in complete M199 media containing 0.05 mM BrdU (Sigma) for 4, 8 or 24 hrs, after which they were washed and fixed in 4% (w/vol) paraformaldehyde in PBS for 20 min and stained as described below. For all BrdU immunodetection experiments, paraformaldehyde-fixed samples were permeabilized with block perm buffer (PBS with 5% normal goat sera and 0.1% Triton-X-100) for 15 min, washed in distilled water, and then immersed in 2M HCl for 40 min. After extensive washing with PBS, the samples were incubated in a block perm buffer for 15 min. Subsequently, BrdU and rGFP were labelled simultaneously overnight at 4 °C by using 1/100 dilution of the rat monoclonal anti BrdU (Abcam) and 1/100 dilution of the rabbit monoclonal anti GFP (Abcam) as primary antibodies. The next day the samples were washed two times with PBS, after which they were treated with block perm buffer for 5 min. Samples were subsequently incubated for 1.5 hrs with Alexa 555 goat polyclonal anti rat IgG (1/1000) and Alexa 488 goat polyclonal anti rabbit IgG (1/1000) (Thermo fisher scientific). After two washes with PBS the samples were mounted with Prolong Gold antifade (Thermo fisher scientific) for confocal microscopy or diluted in PBS for the analysis with the flow cytometer. Microscopy was performed on a Zeiss LSM 700 laser scanning microscope. Image analysis was performed using ImageJ software. The samples for flow cytometry were analyzed with the BD FACS Verse™ in the medium flow rate mode. The time of generation was estimated by extrapolating the time needed to have 100% of the population BrdU^+^ from the percentage of BrdU^+^ cells after 4 hours of incubation.

### Infection in animal models with *L. braziliensis* and *L. mexicana* GFP lines and screening of living amastigotes *ex vivo*

Twelve mice (BALB/c) or hamsters (LVG Golden Syrian Hamster) provided by Charles River were anesthetized by intraperitoneal injection of Ketamine (1mg/10g body weight) and Xylazine (0.08 mg/10g body weight). Anesthetized animals were infected in both hind footpads by sub-cutaneous injection of 5 ×10^5^ stationary phase parasites pre-stained with CellTracker Deep Red in 20 μl of saline solution (0.3 mL syringe and a 30G needle gauge). Four mice and hamsters were sacrificed at 2 weeks, 8 weeks and 16 weeks post infection.

The feet of animals were collected and minced with scissors. The pieces were placed in 1.0 ml of complete M199 containing 200 μg/ml of Liberase TL (Roche) and incubated at 34 °C for 30 min. Subsequently, the samples were filtered with a 70 μm cell strainer and washed with 10 ml of PBS. After centrifugation (1500 g) the pellets were resuspended in 1 mL of complete M199 and aliquot was placed in a 96 well Glass bottom microplates. The screening of living amastigotes was performed with the confocal microscope with settings for the codetection of GFP and the Celltracker Deep Red.

## ACKNOWLEDGEMENTS

We thank Dr. Stephen Beverley and Dr. Michael Mandell for sharing their protocols for the BrdU assay in *Leishmania*.

## SUPPLEMENTAL MATERIAL

**Fig S1.** Live cell imaging of *L. mexicana* Bel21 rGFP Cl1. (**A**) The expression of rGFP was monitored by confocal microscopy in Pro^Log^, Pro^Sta^, Ama^Axe^ and Ama^Int^. As in *L. braziliensis*, a deep downregulation of rGFP was observed in Ama^Axe^ and Ama^Int^ of *L. mexicana*. The scale bar in each figure represents 5 μM. **B**, Quantification of the expression of rGFP over the cell cycle of *L. mexicana* by Flow cytometry. The gray peak in each panel is an overlay of the Wild Type strain (Non GFP negative control). Each histogram represents 10^5^ live cells, all negative for the NucRed staining.

**Fig S2.** Evaluation of the proliferative capability of *L. mexicana* in promastigotes and amastigotes. **A**, Immunodetection (confocal microscopy) of rGFP and BrdU, 8 hrs after exposure with BrdU in Pro^Log^, Pro^Sta^ and Ama^Axe^. **B**, Immuno-detection of BrdU and rGFP in Ama^Int^. In order to be able to monitor Ama^Int^ inside the parasitophorous vacuole the parasites were pre-stained with Cell Tracker before the infection. The scale bar in each picture represents 5 μM.

